# In vitro reconstitution of minimal human centrosomes

**DOI:** 10.1101/2025.02.20.639226

**Authors:** Manolo U. Rios, Weronika E. Stachera, Nicole E. Familiari, Claudia Brito, Thomas Surrey, Jeffrey B. Woodruff

## Abstract

CDK5RAP2/CEP215 is a key pericentriolar material (PCM) protein that recruits microtubule-nucleating factors at human centrosomes. Using an *in vitro* reconstitution system, we show that CDK5RAP2 is sufficient to form micron-scale scaffolds around a nanometer-scale nucleator in a PLK-1-regulated manner. CDK5RAP2 assemblies recruited and activated gamma tubulin ring complexes (γ-TuRCs) which, in the presence of α/β tubulin, generated microtubule asters. We found that F75 in CDK5RAP2 is partially needed to recruit γ-TuRC yet is indispensable for γ-TuRC activation. Furthermore, our system recapitulated key features of centrosome-amplified cancer cells. CDK5RAP2 scaffolds selectively recruited the molecular motor KifC1/HSET, which enhanced concentration of α/β tubulin, microtubule polymerization, and clustering of the assemblies. Our results highlight the specificity and selectivity of *in vitro* generated CDK5RAP2 scaffolds and identify a minimal set of components required for human centrosome assembly and function. This minimal centrosome model offers a powerful tool for studying centrosome biology and dysfunction in human health and disease.

## INTRODUCTION

Centrosomes are major microtubule-organizing centers (MTOCs) in animal cells. These membraneless organelles play a crucial role in mitotic cell division and development (Conduit et al., 2015; Pimenta-Marques and Bettencourt-Dias, 2020). Centrosomes form through structured centrioles that organize a micron-scale layer of protein called pericentriolar material (PCM) that nucleates and anchors microtubules. One of the main proteins underlying PCM formation is CDK5 regulatory subunit-associated protein 2 (CDK5RAP2), also known as CEP215 (Ching et al., 2000; Fong et al., 2008). Dysfunction of CDK5RAP2 can lead to genomic instability, infertility, microcephaly and other developmental disorders (He et al., 2024; Megraw et al., 2011; Nasser et al., 2020; Rahimian et al., 2023; Yigit et al., 2015). Currently, there are several major questions surrounding the molecular mechanisms of CDK5RAP2 assembly and function. First, what are the molecular principles driving CDK5RAP2 accumulation around centrioles? Second, how does CDK5RAP2 bind and activate microtubule nucleating complexes? Third, does this protein serve as a major PCM scaffold? These questions remain incompletely answered due to the lack of a biochemical reconstitution system. Therefore, we set out to generate minimal human centrosomes using purified CDK5RAP2 and other PCM components.

Structural studies of CDK5RAP2 and its functional homologues in flies and worms (Cnn and SPD-5, respectively) show that centrosome assembly is typically achieved in two general steps. First, these proteins are nucleated around the centrioles to form the interphase PCM (Cabral et al., 2019; Fu and Glover, 2012; Hamill et al., 2002; Lawo et al., 2012; Mennella et al., 2012). In humans, nucleation of CDK5RAP2 around the centrioles is mediated by the adaptor protein Pericentrin (Buchman et al., 2010; Wang et al., 2010). Pericentrin decorates the outer wall of the centrioles and binds the conserved CM2 domain in CDK5RAP2 located at its C-terminus (a.a.1680-1893) (Buchman et al., 2010; Kim and Rhee, 2014; Lawo et al., 2012). However, recent evidence shows that CDK5RAP2 can assemble into micron-scale scaffolds in the absence of centrioles (Chen et al., 2022; Watanabe et al., 2020). This suggests that centrioles are not strictly required to nucleate CDK5RAP2 assembly. The universal requirements for PCM nucleation thus require further investigation.

The second major step in centrosome assembly involves increased accumulation of CDK5RAP2 or its functional homologues in preparation for mitosis, also termed centrosome expansion. This process is regulated by mitotic kinases such as Polo-like kinase 1 (PLK-1); in flies and worms, PLK-1 phosphorylation sites on major PCM proteins are known (Conduit et al., 2014; Decker et al., 2011; Dobbelaere et al., 2008; Feng et al., 2017; Sunkel and Glover, 1988; Woodruff et al., 2015). In *D. melanogaster*, centrioles help generate local pulses of PLK-1 activity to initiate PCM expansion (Feng et al., 2017; Wong et al., 2022). Our group and others demonstrated that PLK-1 phosphorylation induces conformational changes in SPD-5 that promotes its multimerization in the presence or absence of centrioles (Mittasch et al., 2020; Ohta et al., 2021; Rios et al., 2024; Woodruff et al., 2015; Wueseke et al., 2016). Human PLK-1 is also necessary for centrosome expansion in human cells (Haren et al., 2009; Santamaria et al., 2011). However, 1) a comprehensive list of PLK-1 phosphorylation sites in CDK5RAP2 and identification of which of these may drive centrosome expansion does not currently exist and 2) it is not known if CDK5RAP2 can multimerize in a PLK-1-dependent fashion without centrioles.

CDK5RAP2 has been proposed to serve as a major PCM scaffold due to its ability to form a stable, salt-resistant network within centrosomes and recruit numerous centrosome-associated proteins (Andersen et al., 2003; Buchman et al., 2010; Ching et al., 2000; Fong et al., 2008; Fong et al., 2009; Graser et al., 2007). CDK5RAP2 plays a pivotal role in recruiting γ-tubulin ring complexes (γ-TuRCs) to the centrosomes (Choi et al., 2010; Fong et al., 2008). γ-TuRCs bind α/β tubulin and promote microtubule nucleation and anchoring (Brito et al., 2024), thus contributing the centrosome’s main function as a MTOC. γ-TuRC anchoring is thought to be mediated by the conserved CM1 domain of CDK5RAP2 located at its N-terminus (a.a.51-100)(Choi et al., 2010). This interaction allows centrosomes to template microtubule asters (Rai et al., 2024; Serna et al., 2024). Phenylalanine 75 (F75) in CDK5RAP2’s CM1 domain is particularly important for its association with γ-TuRCs (Choi et al., 2010). However, it remains unclear if this residue is responsible for γ-TuRC recruitment to the centrosomes and activation.

In addition to γ-TuRCs, CDK5RAP2 interacts with the minus-end directed motor HSET/KifC1 (Chavali et al., 2016). In a wild-type context, HSET is required to keep centrosomes attached to the mitotic spindle poles, while in centrosome-amplified cancer cells, HSET is necessary to cluster supernumerary centrosomes into pseudo-bipolar spindles and promote cancer cell survival (Chavali et al., 2016). Other studies have shown that HSET is sufficient to cluster microtubules and that IFT proteins, in complex with HSET, promote clustering (Hentrich and Surrey, 2010; Norris et al., 2018; Roostalu et al., 2018; Vitre et al., 2020). However, the minimal components sufficient to achieve centrosome clustering remain unclear.

To deepen our understanding of CDK5RAP2 nucleation, supramolecular scaffold assembly and regulation, and microtubule organizing function, we developed a novel *in vitro* reconstitution system using purified human proteins. We found that purified CDK5RAP2 self-assembles into micron-scale scaffolds in the presence of crowding agents or when recognized by a pentameric antibody (IgM) targeting the CM2 domain. This system offers a robust assay to explore CDK5RAP2’s regulatory interactions, as the CDK5RAP2 assemblies can be modulated by human PLK-1, which enhances their size. Using proteomics, we mapped PLK-1 phosphorylation sites in full-length CDK5RAP2. Additionally, the reconstituted CDK5RAP2 assemblies were capable of nucleating microtubule asters in the presence of purified α/β tubulin and human γ-TuRCs. The specificity and selectivity of this feature was validated using a CDK5RAP2(F75A) mutant, revealing that CDK5RAP2 macromolecular scaffolds by themselves cannot bind α/β tubulin. *In vitro*-generated CDK5RAP2 scaffolds also selectively recruited wild-type HSET, but not a mutant lacking its N-terminus intrinsically disordered region (IDR). Our results support HSET’s functional relevance in clustering centrosomes and organizing microtubule arrays *in vitro*.

## RESULTS

### Purified human CDK5RAP2 can assemble into micron-scale scaffolds *in vitro*

In mitotic human cells, CDK5RAP2 molecules accumulate around centrioles to build a spherical structure 1-2 μm in diameter (**Fig.1A**)(Fong et al., 2008; Lawo et al., 2012). CDK5RAP2 is expressed as multiple isoforms. AlphaFold2 predicts that the longest isoform encodes a 215 kDa protein comprising 19 alpha-helical regions interspersed by intrinsically disordered linkers (Jumper et al., 2021; Varadi et al., 2021; Varadi et al., 2023). CDK5RAP2 isoform B (Protein ID Q96SN8-4, lacking a.a. 1576-1654) is a dominant splice variant expressed in HeLa and colon cancer cells (Harrison et al., 2018; He et al., 2024; Hein et al., 2015). To investigate if CDK5RAP2 can self-assemble into a multimeric scaffold, we first expressed and purified GFP-tagged CDK5RAP2 isoform B from SF9 cells (**Fig. 1A**). At near-physiological salts and protein concentrations (10-120 nM protein, 150 mM KCl;(Hein et al., 2015)) GFP::CDK5RAP2 was soluble and did not form micron scale scaffolds (**Fig. S1A**). We hypothesized that to multimerize *in vitro*, CDK5RAP2 may require conditions that more closely resemble the intracellular environment. First, we mimicked the crowded environment inside living cells using molecular crowding agents. Using as little as 3% (w/v) polyethylene glycol 3000 (PEG), GFP::CDK5RAP2 formed micron-scale assemblies *in vitro* (**Fig.S1A, B**). The size and number of these assemblies scaled positively with the concentration of CDK5RAP2 (**Fig. 1B, C**) and PEG (**Fig. S1B**). Other crowding agents (lysozyme, polyvinylpyrrolidone (PVP), Dextran, and Ficoll) also induced CDK5RAP2 self-assembly (**Fig. S1C, D**). Thus, macromolecular crowding, and not PEG *per se*, is sufficient to promote CDK5RAP2 assembly into micron-scale structures.

**FIGURE 1.**
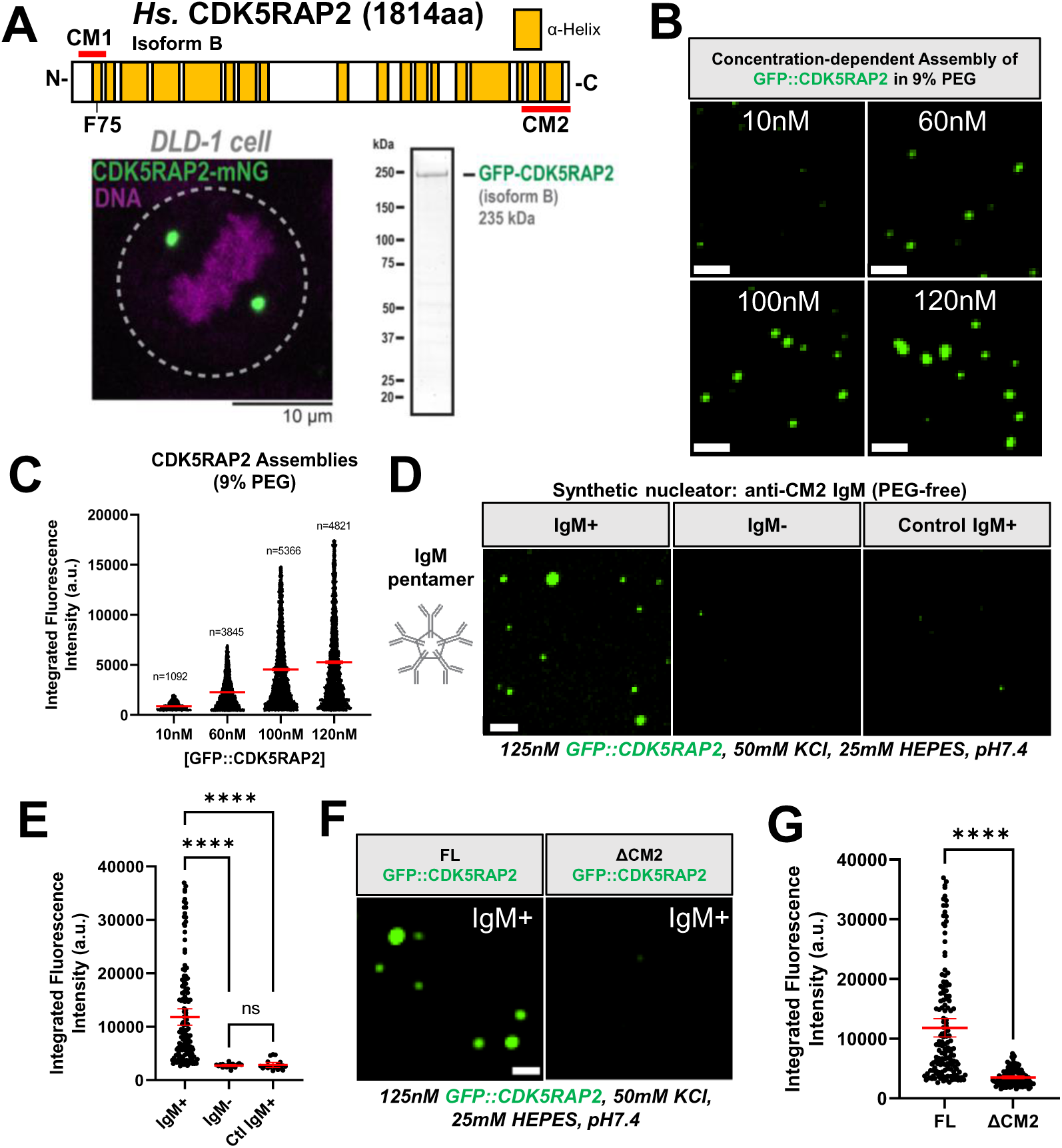
Purified human CDK5RAP2 is sufficient to form micron scale scaffolds. A) Linear representation of *Homo Sapiens* (*Hs.*) CDK5RAP2/CEP215 Isoform B, human colon cancer DLD-1 cell expressing CDK5RAP2-mNeonGreen (mNG) and purified GFP::CDK5RAP2 isoform B from SF9 insect cells. Yellow blocks in linear CDK5RAP2 diagram represent alpha-helical regions predicted by AlphaFold2. CM1 (a.a. 51-100) and CM2 (a.a. 1715-1814) domains are indicated by red bars. The CM1 domain contains F75. B) Purified GFP::CDK5RAP2 combined with 9% PEG at various concentrations (10nM, 60nM, 100nM and 120nM). Scale bar, 5 μm. C) Quantification of panel 1B (mean +/- 95% C.I.; 10nM (n=1092 assemblies), 60nM (n=3845 assemblies), 100nM (n=5366 assemblies), 120nM (n=4821 assemblies). D) GFP::CDK5RAP2 assemblies were nucleated by adding <11 pM IgM raised against the CM2 domain of CDK5RAP2 (IgM+). Addition of buffer with no IgM (IgM-) or an anti-myosin heavy chain (Control IgM) did not nucleate micron-scale GFP::CDK5RAP2 assemblies. Scale bar, 5 μm. E) Quantification of panel 1D (mean +/- 95% C.I.; IgM+ (n=166 assemblies), IgM- (n=23 assemblies), Ctl IgM (n=20 assemblies)). P values from One-way ANOVA followed by Tukey’s multiple comparisons test. F) Assembly of full length (FL) GFP::CDK5RAP2 into micron-scale scaffolds (left panel). GFP::CDK5RAP2 lacking the CM2 domain does not form micron-scale assemblies (right panel). This domain (last 200 a.a.) is recognized by the IgM. Scale bar, 5 μm. G) Quantification of panel 1F. All data points represent individual GFP::CDK5RAP2 assemblies (mean +/- 95% C.I.; (FL (n=166 assemblies), ΔCM2 (n=131 assemblies)) P value from a student t-test.

Although effective, assembly of CDK5RAP2 into micron-scale scaffolds using synthetic crowders may not recapitulate regulated assembly. Thus, we devised an alternative way to generate CDK5RAP2 micron-scale assemblies. CDK5RAP2 naturally docks with the centriole indirectly via its CM2 domain (Wang et al., 2010). Thus, we tested the effect of introducing a pentameric IgM antibody (valence = 10 binding sites) raised against the CM2 domain of CDK5RAP2. We hypothesized that by using an antibody that closely recapitulates the geometry, valence, and scale of centrioles we could mimic centriolar nucleation and lower the energy barrier for CDK5RAP2 assembly. Indeed, in the presence of less than 11 pM anti-CM2 CDK5RAP2 IgM, we observed spherical, micron-scale GFP::CDK5RAP2 assemblies (**Fig. 1D, E**, **Table S1**). No assemblies formed when IgM was absent or when a control anti-myosin heavy chain IgM was used (**Fig. 1D, E**). For the antibody to produce robust assemblies, it was essential to reduce the amount of KCl to 50 mM. To further test the specificity and assembly capabilities of the CDK5RAP2 antibody we compared the assembly of GFP::CDK5RAP2 lacking its CM2 domain (ΔCM2) with full-length protein. Only full-length GFP::CDK5RAP2 formed robust micron-scale assemblies in the presence of the IgM nucleator (**Fig. 1F, G**). Overall, our observations demonstrate the ability of CDK5RAP2 to multimerize in the presence of crowding agents or a synthetic nucleator. Furthermore, our results reinforce *in vivo* findings demonstrating that centrioles are not strictly required for CDK5RAP2 micron-scale assembly (Chen et al., 2022).

### PLK-1 phosphorylation increases the size of CDK5RAP2 scaffolds *in vitro*

In human cells, PLK-1 kinase activity is required to expand the amount of CDK5RAP2 at centrosomes ∼5-fold during mitosis (Haren et al., 2009; Santamaria et al., 2011). We thus tested if PLK-1 is sufficient to regulate the assembly of our antibody-nucleated GFP::CDK5RAP2 assemblies. We incubated pre-dephosphorylated full-length GFP::CDK5RAP2 + anti-CM2 IgM in the presence of purified kinase dead (KD) or constitutively active (CA) human PLK-1 and ATP·MgCl_2_ for 1 hr at 23°C. Confocal microscopy revealed that PLK-1(CA) increased the mass of GFP::CDK5RAP2 assemblies 1.7-fold compared to the assemblies incubated with PLK-1(KD)(P<0.0001) (**Fig. 2A, B**). This suggests our system is subject to physiological regulation and that PLK- 1 promotes centrosome assembly by directly enhancing CDK5RAP2 self-assembly. Our data also suggest that additional factors are required to achieve full-scale PCM growth seen in cells.

**FIGURE 2.**
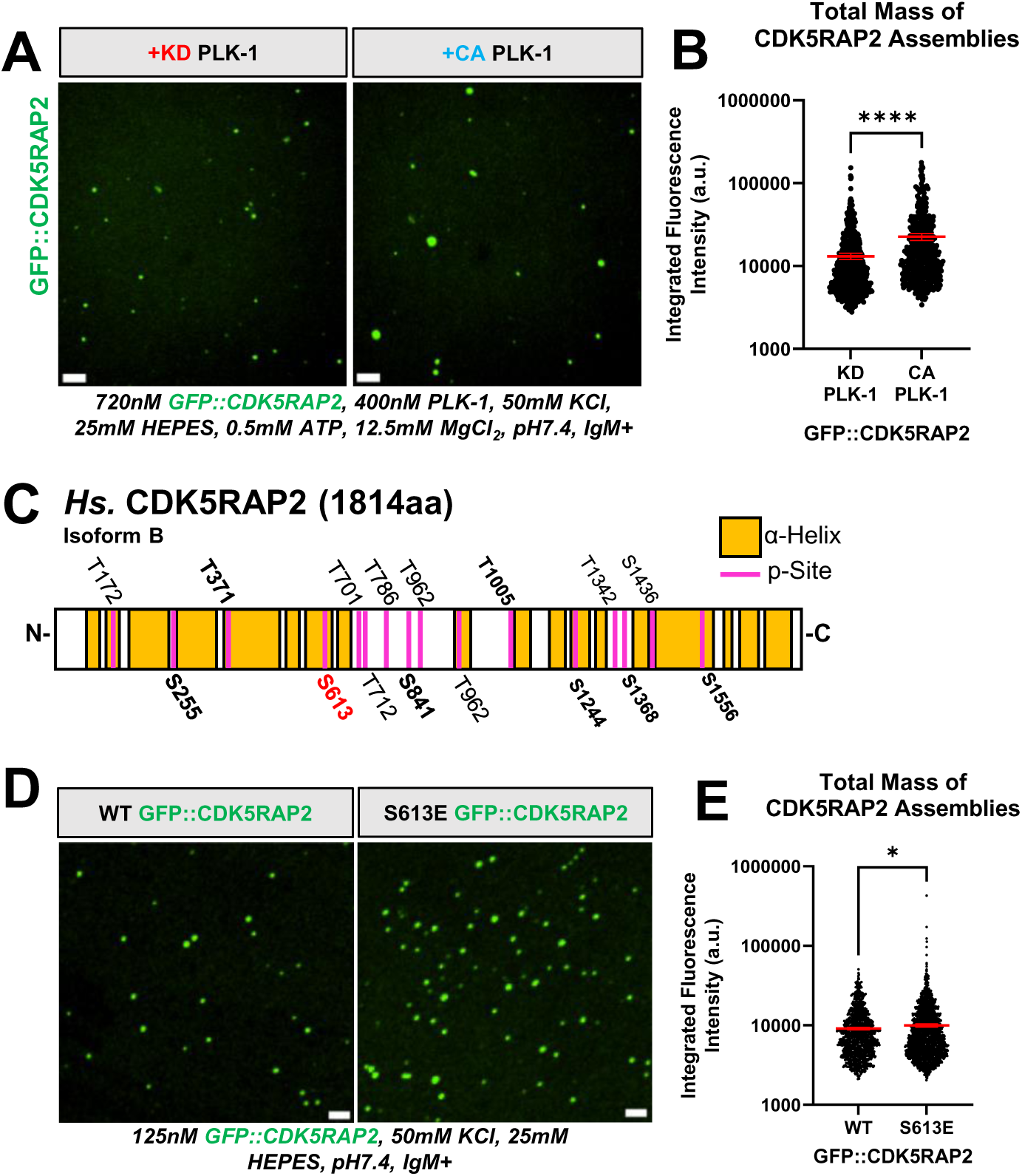
Phospho-regulation of *in vitro* CDK5RAP2 scaffolds. A) GFP::CDK5RAP2 assemblies generated in the presence of purified kinase dead (KD) or constitutively active (CA) human PLK-1. Scale bar, 5 μm. B) Quantification of panel 2A. Data points represent individual GFP::CDK5RAP2 assemblies (mean +/- 95% C.I.; KD PLK-1 condition (n=604 assemblies), CA PLK-1 condition (n=511 assemblies)). Significant differences were assessed using a student t-test. C) Identified PLK-1 phosphorylation sites (p-Sites) in GFP::CDK5RAP2 incubated with human PLK-1 plus ATP·MgCl2. Bold p-sites are also found in PhosphoSite Plus. S613 (red) is phosphorylated by PLK-1 *in vivo* and *in vitro*. Control reactions consisted of pre-dephosphorylated GFP::CDK5RAP2 plus PLK-1 but no ATP·MgCl2. D) *In vitro* assemblies of pre-dephosphorylated GFP::CDK5RA2 and pre-dephosphorylated GFP::CDK5RAP2(S613E). Scale bar, 5 μm. E) Quantification of total mass of CDK5RAP2 assemblies in panel D. Each data point represents individual GFP::CDK5RAP2 assemblies (mean +/- 95% C.I.; WT (n=1337 assemblies), S613E (n=2463 assemblies)). Significant differences were assessed using a student t-test.

To determine which phospho-residues in CDK5RAP2 may be involved in its assembly, we mapped phosphorylation sites using mass spectrometry. We identified 16 phosphorylated residues in CDK5RAP2, 56% of which fell within disordered linker regions, while the remaining 44% were in alpha helical regions (**Fig. 2C**). These phospho- sites were found in samples containing PLK-1 but not in samples containing only pre-dephosphorylated CDK5RAP2. We also compared the phosphorylation sites found in our study to annotated phosphorylation sites of human CDK5RAP2 found in the PhosphoSite Plus (PSP) database (Hornbeck et al., 2015). Eight of the PSP-annotated residues were detected in our experiment: S255, T371, S613, S841, T1005, S1244, S1368, and S1556 (**Fig. 2C**, **S2A**, **B**). Of these, S613 is the only residue that has been verified to be phosphorylated by PLK-1 *in vivo* (Santamaria et al., 2011). The kinase or kinases responsible for the phosphorylation of the other previously reported residues in the PSP were unidentified; our results suggest they may be PLK-1 sites. Our experiment revealed 8 novel PLK-1 phosphorylation sites in CDK5RAP2: T172, T701, T712, T786, S872, T962, T1342, and S1436 (**Fig. 2C**). We did not observe residues known to be phosphorylated by the mitotic kinase LARRK1 (pS102 or pS140) (Hanafusa et al., 2015), indicating that our experimental conditions allow for kinase specificity.

Given that S613 was the only residue unambiguously identified to be phosphorylated by PLK-1 *in vivo* and *in vitro,* we tested if phosphorylation of this single residue is sufficient to enhance CDK5RAP2 self-assembly *in vitro*. We purified pre-dephosphorylated GFP::CDK5RAP2 containing a glutamic acid substitution at S613 (phospho-mimetic) and compared its self-assembly capabilities to pre-dephosphorylated “wild-type” GFP::CDK5RAP2. The reaction contained the anti-CM2 antibody but no PLK-1 or ATP·MgCl_2_. GFP::CDK5RAP2(S613E) assemblies were, on average, 1.1-fold more massive than WT assemblies (P<0.02)(**Fig. 2D, E**). This result suggests that phosphorylation of residue S613 weakly enhances growth of CDK5RAP2 assemblies. Since this does not reach the effect of adding PLK-1(CA), multiple additional phosphorylation sites likely contribute to full-scale self-assembly of CDK5RAP2. We conclude that *in vitro* minimal centrosome scaffolds are subject to phospho-regulation by PLK-1, a key feature of *in vivo* PCM. Our results suggest that phosphorylation of CDK5RAP2 promotes its multimerization in the absence of centrioles, similar to the mechanism proposed for SPD-5 (*C. elegans*) (Rios et al., 2024; Woodruff et al., 2015). However, centrioles, other inner PCM proteins, and synergistic cooperation between multiple phosphorylated residues are likely critical to maintain PLK-1 activity and achieve maximal PCM size (Conduit et al., 2014; Rios et al., 2024; Wong et al., 2022; Woodruff et al., 2015).

### *In vitro* CDK5RAP2 micron-scale assemblies selectively recruit γ-TuRCs and nucleate microtubule asters

*In vivo*, CDK5RAP2 recruits γ-TuRCs to produce robust microtubule asters, giving centrosomes their main function (Choi et al., 2010; Fong et al., 2008). However, it is not known if CDK5RAP2 and γ-TuRCs are sufficient to achieve robust microtubule aster formation or if additional components are required. We first used confocal microscopy to test if CDK5RAP2 assemblies nucleate microtubules by themselves by assembling them with anti-CM2 IgM and HiLyte-647-labeled α/β tubulin. At 37°C, GFP::CDK5RAP2 assemblies alone did not recruit α/β tubulin or generate microtubule asters (**Fig. S3A**).

We then added purified human γ-TuRCs tagged with blue fluorescent protein (mBFP). CDK5RAP2 assemblies recruited γ-TuRCs and α/β tubulin (**Fig. 3A-C**). There was a strong correlation between the amount of γ-TuRC fluorescence and α/β tubulin recruited (Pearson’s correlation r=0.86; P<0.0001) (**Fig. S3B**). 90% of γ-TuRC-positive assemblies formed microtubule asters over the course of 30 min (n = 134 assemblies) (**Fig. 3A-D**, **Supplemental Movie 1**). Thus, a minimal module of self-assembled CDK5RAP2, γ- TuRCs, and α/β tubulin is sufficient to recapitulate microtubule aster formation, a key feature of centrosome function in cells.

**FIGURE 3.**
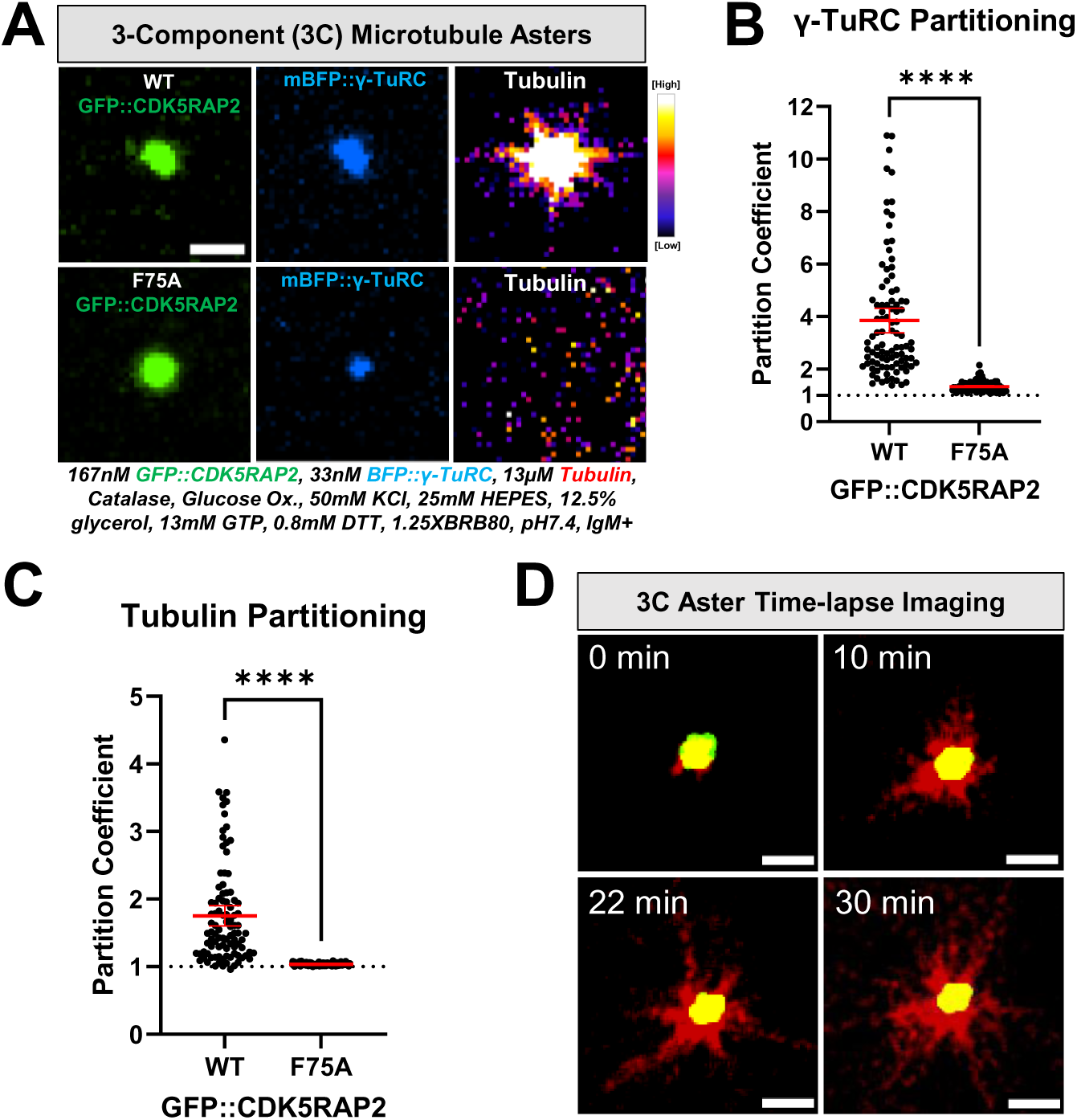
Microtubule nucleating activity of CDK5RAP2 scaffolds with γ-TuRCs A) 3-Component (3C) reactions were assembled with purified GFP::CDK5RAP2 (WT or F75A), mBFP::γ-TuRCs, and HiLyte647-labeled α/β tubulin. Scale bar, 3μm B) Quantification of γ-TuRC partitioning into GFP::CDK5RAP2 assemblies (mean +/- 95% C.I.; WT, n=95; F75A, n=117) . Significant differences were assessed using a student t-test. C) Quantification of α/β tubulin partitioning into GFP::CDK5RAP2 assemblies (mean +/- 95% C.I.; WT, n=95; F75A, n=117). Significant differences were assessed using a student t-test. D) Time-lapse imaging of microtubule growth from GFP::CDK5RAP2 scaffolds + mBFP::γ-TuRCs. Z-projections were obtained every 2 min for 30 min at 37°C. Scale bar, 3μm.

CDK5RAP2 binds γ-TuRCs through its N-terminal CM1 domain (Choi et al., 2010; Fong et al., 2008). Mutation of phenylalanine 75 to alanine (F75A) in a truncated CDK5RAP2 construct (a.a. 51-100) disrupts γ-TuRC binding and subsequent ring closure (Brito et al., 2024; Choi et al., 2010; Xu et al., 2024). Whether F75 is critical for binding or activation of γ-TuRC in the context of full-length CDK5RAP2 has not been tested. Purified CDK5RAP2(F75A) multimerized into micron-scale assemblies that poorly recruited γ- TuRCs and α/β tubulin (**Fig. 3A-C**). This result reinforces the importance of F75 for the interaction between γ-TuRC and CDK5RAP2 and demonstrates that our system recapitulates binding specificity. The fact that the F75A mutant still recruits minute amounts of γ-TuRC suggests that additional interaction motifs besides CM1 could engage with the γ-TuRC. In these mutant assemblies, there was no correlation between γ-TuRC and α/β tubulin recruitment, indicating that the bound γ-TuRCs were incompetent to bind α/β tubulin, unlike the control case (**Fig. S3C**). Consistently, these assemblies did not nucleate microtubule asters (0%, n = 490 assemblies) (**Fig. 3A**). Thus, F75 is required for activating the γ-TuRC by improving its ability to bind α/β tubulin. We propose that microtubule aster formation involves a two-step process of γ-TuRC binding to CDK5RAP2 and subsequent activation via interactions mediated by F75.

### Kinesin HSET/KifC1 potentiates microtubule aster formation and induces clustering of CDK5RAP2 assemblies *in vitro*

Vertebrate centrosomes stay connected to the mitotic spindle in part through the activity of HSET/KifC1, a kinesin-14 motor protein that directly binds microtubules and CDK5RAP2 (Chavali et al., 2016; Mountain et al., 1999). In cancer cells with amplified centrosomes, HSET plays a critical survival role by clustering centrosomes to prevent multipolar spindle formation (Chavali et al., 2016). We tested if our *in vitro* system could be used as tool to investigate how HSET contributes to centrosome-spindle attachment and centrosome clustering. Thus, we first probed the ability of CDK5RAP2 to recruit HSET. We purified full-length mCherry-tagged HSET (WT), an HSET mutant that cannot bind CDK5RAP2 in cells (ΔIDR; missing a.a. 2-138 (Chavali et al., 2016)) and mCherry alone as a control. We then assessed their partitioning into GFP::CDK5RAP2 assemblies *in vitro*. mCherry::HSET(WT) directly interacted with GFP::CDK5RAP2 scaffolds whereas mCherry::HSET(ΔIDR) and mCherry alone did not (**Fig. 4A, B**). This result further demonstrates that our *in vitro* CDK5RAP2 scaffolds are specific in their recruitment of clients, recapitulating a key functional aspect of *in vivo* PCM.

**FIGURE 4.**
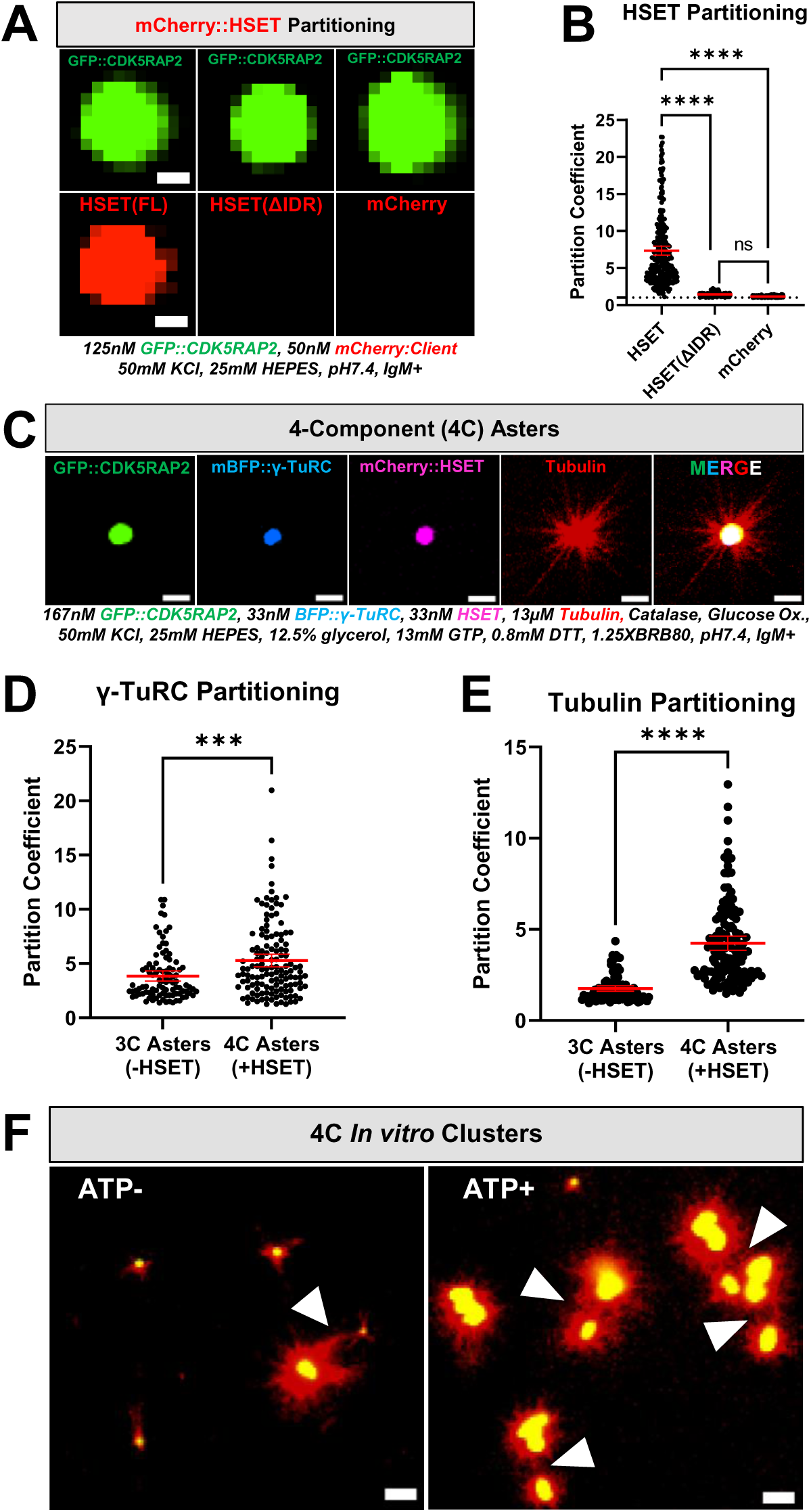
Selective HSET recruitment, microtubule-nucleating effects, and clustering of CDK5RAP2 scaffolds *in vitro*. A) Selective recruitment of purified mCherry::HSET (full-length), mCherry::HSET lacking its intrinsically disordered region (τιIDR, missing a.a. 2-138), or mCherry into micron-scale GFP::CDK5RAP2 assemblies. Scale bar, 1µm. B) Quantification of panel 4A (mean +/- 95% C.I.; HSET(FL), n=293 scaffolds; HSET(ΔIDR), n=294 scaffolds; mCherry, n=285 scaffolds. Each data point represents a CDK5RAP2 scaffold. Significant differences were assessed using One-way ANOVA followed by Tukey’s multiple comparisons test. HSET(FL) n=293 scaffolds, HSET(ΔIDR) n=294 scaffolds, mCherry n=285 scaffolds. C) Z-projection of 4-Component (4C) asters consisting of 167nm GFP::CDK5RAP2, 33nM mBFP::γ-TuRCs, 33nM mCherry::HSET and 13µM HiLyte-647-labeled α/β tubulin. Buffer consists of 50mM KCl, 25mM HEPES, pH7.4, IgM+. D) Quantification of γ-TuRC partitioning in 3-component (3C) vs 4-component (4C) asters (mean +/- 95% C.I.; 3C, n=95; 4C, n=133). Significant differences were assessed using a student t-test. E) Quantification of α/β tubulin partitioning in 3-component (3C) vs 4-component (4C) asters. (mean +/- 95% C.I.; 3C, n=95; 4C, n=133). Significant differences were assessed using a student t-test. F) Z-projection of clusters formed by 4C asters in the presence of 0 or 0.66mM ATP plus MgCl_2_. Arrowheads indicate microtubule contacts between GFP::CDK5RAP2 scaffolds. Scale bar, 5µm.

Next, we asked if *in vitro*-generated CDK5RAP2 scaffolds can bind multiple clients simultaneously and if HSET is sufficient to cluster aster-forming CDK5RAP2 assemblies *in vitro*. Thus, we assembled GFP::CDK5RAP2 scaffolds using the anti-CM2 IgM along with human mCherry::HSET and mBFP::γ-TuRCs. Then, we combined these assemblies with HiLyte647-labeled α/β tubulin and no ATP·MgCl_2_. All proteins co-localized to the micron-scale GFP::CDK5RAP2 scaffolds simultaneously to generate robust microtubule asters (**Fig. 4C**). We conclude that CDK5RAP2 can serve as a major centrosome scaffold for multiple clients *in vitro*.

In the presence of mCherry::HSET, GFP::CDK5RAP2 scaffolds recruited 1.4-fold more γ-TuRCs and 2.4-fold more α/β tubulin (**Fig. 4D, E**). More α/β tubulin partitioned into HSET-containing CDK5RAP2 scaffolds, even when similar amounts of γ-TuRCs were recruited (**Fig. S4A**). In addition, HSET could recruit α/β tubulin to scaffolds lacking γ- TuRCs in the presence of ATP_●_MgCl_2_ but could not make asters (**Fig. S4B**). We conclude that HSET recruits α/β tubulin into CDK5RAP2 assemblies without needing γ-TuRC or ATP. We hypothesize HSET can potentiate CDK5RAP2/γ-TuRC aster-formation capabilities through increased α/β tubulin and γ-TuRC partitioning and perhaps via γ- TuRC activation.

Using static Z-projections and time-lapse recordings, we investigated if these 4- component asters could cluster together in the presence or absence of ATP·MgCl_2_ on passivated glass slides. Under these conditions, CDK5RAP2 scaffolds readily formed clusters bridged by microtubules in the presence of ATP (25 clusters, n=5 images), but less so in the absence of ATP (8 clusters, n=5 images) (**Fig. 4F**, **S4C**). Once the microtubule bridges had formed, we did not observe further motor-mediated constriction of CDK5RAP2 assemblies (**Supplemental Movie 2**). We did not detect significant differences in area and aspect ratio of CDK5RAP2 assemblies between these conditions, suggesting that HSET motor activity does not affect CDK5RAP2 self-assembly (**Fig. S4D, E**). These results demonstrate that our system recapitulates HSET binding specificity and centrosome clustering *in vitro.* We conclude that HSET possesses two activities within CDK5RAP2 scaffolds: 1) an ATP-independent ability to concentrate tubulin and γ-TuRC and 2) an ATP-dependent ability to cluster them via microtubules.

## DISCUSSION

The outer layer of centrosomes, called PCM, is a dynamic supramolecular scaffold that organizes a radial array of microtubules for mitotic cell divisions. Our results reveal a minimal set of components sufficient to recapitulate human PCM assembly, microtubule aster formation, and ATP-dependent clustering. These components include a coiled-coil rich scaffold protein (CDK5RAP2), a microtubule nucleating complex (γ-TuRC), an ATP- powered motor protein (HSET), and α/β tubulin. In addition to these native proteins, specific conditions are needed to overcome the energy barrier for scaffold assembly, including macromolecular crowding or a nucleation surface (e.g., a pentameric IgM that recruits CDK5RAP2). This system also revealed that the mitotic kinase PLK-1 can directly phosphorylate CDK5RAP2 and enhance its assembly into micron-scale scaffolds. Specific mutations that were identified to disrupt centrosome function *in vivo*— CDK5RAP2(F75A), HSET(τιIDR)—disrupted γ-TuRC activation and HSET recruitment to CDK5RAP2 scaffolds *in vitro.* Finally, our system revealed a new, ATP-independent role for HSET in concentrating α/β tubulin and enhancing microtubule aster formation within the CDK5RAP2 assemblies. Thus, our reconstitution system recapitulates key features of *in vivo* PCM—including specificity of protein recruitment, regulation, and activity—and reveals new insights that could not be achieved through traditional experiments in cells. Based on these results, we propose the following model for PCM assembly and function in human cells. First, the PCM scaffold assembles through multimerization of CDK5RAP2 around a nucleation point (e.g., the centriole) using a conserved centrosome localization sequence (CM2). Second, PLK-1 phosphorylates CDK5RAP2 to potentiate its multimerization. Third, the PCM scaffold concentrates microtubule nucleating complexes, which subsequently concentrate soluble α/β tubulin and initiate microtubule polymerization.

Prior mechanistic understanding of kinase-driven PCM expansion was largely derived from studies in *Drosophila* and *C. elegans*. In these species, Polo Kinase phosphorylates coiled-coil containing PCM proteins (Cnn and SPD-5, respectively) at multiple sites (Conduit et al., 2014; Woodruff et al., 2015). This activity is thought to release autoinhibitory folding, thus promoting interactions between coiled-coil motifs *in trans* to build a supramolecular scaffold (Feng et al., 2017; Rios et al., 2024). Here, we provide evidence that vertebrate CDK5RAP2 can multimerize in a Polo Kinase-potentiated fashion *in vitro*, suggesting that it is a functional homolog of Cnn and SPD-5. Furthermore, we saw that a single phospho-mimicking point mutation on CDK5RAP2, S613E, had a mild effect on GFP::CDK5RAP2 scaffold nucleation and growth *in vitro*. This effect was not as dramatic as natural PCM growth in cells; thus, we speculate that multiple phosphorylated residues, and likely other proteins, cooperate to achieve full-scale expansion of the human PCM scaffold. Although incomplete, this simple reconstitution system nevertheless provides a strong foundation towards revealing the molecular mechanism of PLK-1-dependent centrosome expansion. Future studies should focus on dissecting this mechanism and testing if PLK-1 phosphorylation induces conformational changes in CDK5RAP2 to promote its self-assembly, both *in vitro* and in cells.

Our experiments also shed insight into microtubule nucleation mechanisms and the minimal requirements for centrosome clustering. Previous experiments revealed that a truncated version of CDK5RAP2 (a.a. 51-100), but not an F75A mutant, was sufficient to pull down γ-TuRC in cell extracts (Choi et al., 2010). This result suggested that CDK5RAP2 recruits γ-TuRC through its CM1 domain. Whether F75 is critical in the context of full-length CDK5RAP2 had not been tested. We saw that assemblies made with either full-length wild-type CDK5RAP2 or the F75A mutant could recruit γ-TuRC, although the mutant showed reduced binding. However, γ-TuRCs were only active when bound to wild-type CDK5RAP2. These data indicate that 1) γ-TuRC binding to CDK5RAP2 and its activation are separate steps, 2) additional domains beyond CM1 mediate γ-TuRC recruitment, and 3) F75 is critical for activating γ-TuRCs, likely due to mediating ring closure (Xu et al., 2024).

Addition of the kinesin-14 motor HSET improved microtubule aster formation by CDK5RAP2/γ-TuRC assemblies. However, the mechanism of this synergy is not fully clear. In the presence of soluble tubulin and without CDK5RAP2, HSET can form microtubule asters using its N-terminal IDR and C-terminal motor domain (Hentrich and Surrey, 2010; Norris et al., 2018; Roostalu et al., 2018). We found that HSET by itself did not form microtubule asters when localized within the CDK5RAP2 scaffold but could recruit α/β tubulin. We speculate that HSET’s intrinsic aster-formation capabilities are hindered by its interaction with the CDK5RAP2 scaffold, presumably due to preferential binding of CDK5RAP2 for HSET’s IDR. HSET also improved γ-TuRC recruitment into CDK5RAP2 scaffolds. Thus, HSET likely promotes microtubule polymerization within the PCM by locally concentrating soluble α/β tubulin and γ-TuRC in an ATP-independent manner. Whether HSET can directly improve the activity of γ-TuRC should be tested in the future. Adding ATP promoted clustering of the CDK5RAP2/γ-TuRC/HSET assemblies, consistent with a reported role for HSET motor activity in clustering multiple centrosomes in cells (Chavali et al., 2016). Based on these observations, we propose that HSET drives centrosome clustering by promoting microtubule polymerization through concentration of soluble tubulin and crosslinking microtubules.

*In vitro* reconstitution systems have been instrumental in elucidating mechanisms of centrosome assembly and function in invertebrates (Conduit et al., 2014; Feng et al., 2017; Rios et al., 2024; Schnackenberg et al., 1998; Woodruff et al., 2017). However, a robust bottom-up system using human proteins has remained elusive until now. Our work provides a versatile tool for studying CDK5RAP2 function and its regulation in isolation from cellular complexity. This system recapitulates important centrosome behaviors: PCM scaffold nucleation, PCM expansion, phospho-regulation, microtubule nucleation and client selectivity. Future improvements include: 1) more robust phospho-regulation of CDK5RAP2 scaffold expansion, perhaps via inclusion of different kinases and phosphatases; 2) inclusion of other potential scaffold proteins such as Cep192 and Pericentrin; and 3) inclusion of other effector molecules such as microtubule nucleating or capping complexes. Nevertheless, in its current state, our system has revealed new mechanistic insight into human centrosome biology and holds promise for investigating centrosome dysfunction in human disease.

## ACKNOWLEDGEMENTS

We thank the UT Southwestern Proteomics Core Facility for phosphor-proteomic analysis and Andrew Holland for providing the DLD-1 cells.

## AUTHOR CONTRIBUTIONS

M. Rios performed all experiments and analyzed data. N. Familiari generated baculovirus stocks. W. Stachera helped with cloning and protein purification. C. Brito and T. Surrey purified human γ-TuRC and provided valuable experimental and manuscript feedback. J. Woodruff and M. Rios wrote the manuscript.

## MATERIALS AND METHODS

### Experimental model and subject details

For the expression of recombinant proteins (listed in **Table S2**), we used SF9- ESF *Spodoptera frugiperda* insect cells (Expression Systems) grown at 27°C in ESF 921 Insect Cell Culture Medium (Expression Systems) supplemented with Fetal Bovine Serum (2% final concentration).

### Protein purification

All expression plasmids are listed in **Table S2**. Full-length human GFP::CDKRAP2 proteins (WT, F75A and ΔCM2), unlabeled full-length CDK5RAP2, human PLK-1s (constitutively active and kinase dead) and mCherry::HSET constructs (full-length, ΔIDR) were inserted into a baculoviral expression plasmid (pOCC27 or pOCC28) using standard restriction cloning. Baculoviruses were generated using the FlexiBAC system (Lemaitre et al., 2019) in SF9 cells. Protein was harvested 72 hr after infection during the P3 production phase. Cells were collected, washed, and resuspended in harvest buffer (25 mM HEPES, pH 7.4, 150 mM NaCl). All subsequent steps were performed at 4°C. Cell pellets were resuspended in Buffer A (25 mM HEPES, pH 7.4, 30 mM imidazole, 500 mM KCl, 0.5 mM DTT, 1% glycerol, 0.1% CHAPS) + protease inhibitors and then lysed using a dounce homogenizer. Proteins were bound to Ni-NTA (Qiagen), washed with 10 column volumes of Buffer A, and eluted with 250 mM imidazole. The CDK5RAP2 eluates were then bound to maltose-binding protein (MBP) trap beads (NanoTag Biotechnologies) and washed with five column volumes of Buffer C (25 mM HEPES, pH 7.4, 500 mM NaCl, 0.5 mM DTT, 1% glycerol, 0.1% CHAPS). Proteins were eluted by adding PreScission protease, incubating overnight, and then passing over Ni-NTA beads to remove the Precission protease. Eluted proteins were then concentrated using 3–10 K MWCO Amicon concentrators (Millipore). All proteins were aliquoted in PCR tubes, flash-frozen in liquid nitrogen, and stored at −80°C. Protein concentration was determined by measuring absorbance at 280 nm using a NanoDrop ND-1000 spectrophotometer (Thermo Fisher Scientific). To dephosphorylate CDK5RAP2 proteins, MBP-trap bound GFP::CDK5RAP2 was incubated for 1 h at room temperature in dephosphorylation buffer (1× PMP buffer (NEB) + 1 mM MnCl_2_), + 40,000 U lambda phosphatase (400,000 U/ml, NEB). Beads were then washed twice with buffer C at 4°C during the regular washing steps (total of 7 washes) to remove lambda phosphatase. Dephosphorylation of GFP::CDK5RAP2 was confirmed by PTM identification using mass spectrometry showing effective removal of 99.9–100% of phosphates.

Human γ-TuRCs were purified as previously described (Brito et al., 2024). In short, HELA-Kyoto cells expressing mBFP::γTuRCs were resuspended in lysis buffer (50 mM HEPES, 150 mM KCl, 5 mM MgCl2, 1 mM EGTA, 1 mM DTT, 0.1 mM GTP, pH 7.4) containing protease inhibitors and DNAse I (10 μg ml-1, Sigma-Aldrich). Resuspended cells were lysed, clarified twice by centrifugation (17,000xg, 15 min, 4°C) and filtered. Lysates were desalted, supplemented with protease inhibitors and loaded onto a 1 mL HiTrap SP Sepharose FF column connected in tandem with 1 mL streptavidin mutein matrix beads (Sigma Aldrich) packed into a Tricorn 5/50 column (GE-Healthcare). The streptavidin mutein matrix column was washed with 30 mL storage buffer, 30 mL wash buffer (lysis buffer containing 200 mM KCl and 0.2% (vol./vol.) Brij-35) and 30 mL storage buffer. Proteins were eluted with storage buffer supplemented with 5 mM D-biotin. The buffer was then exchanged into storage buffer, centrifuged (17,000xg, 10 min, 4°C) and separated by size exclusion chromatography using a Superose 6 10/300 GL column. γTuRC peak fractions were pooled, concentrated, ultracentrifuged (278,088.3xg, 10 min, 4°C), snap frozen and stored in liquid nitrogen. Porcine tubulin was used in this study and was purified as previously described (Gell et al., 2011).

### *In vitro* assembly of CDK5RAP2 scaffolds

Purified GFP::CDK5RAP2 stocks were dialyzed prior to their use in reaction buffer (50mM KCl, 25mM HEPES, pH7.0). Dialysis took place overnight at 4°C using 10K MWCO MINI Dialysis Units (ThermoScientifc). Assembly reactions were carried on in 10μL volumes at room temperature for 1-2 minutes. Final protein concentrations and buffer conditions were 0-120mM GFP-CDK5RAP2, 150mM KCl, 25mM HEPES, pH7.4 for crowder-based assemblies and 125mM GFP-CDK5RAP2, 50mM KCl, 25mM HEPES, pH7.4 for antibody-based assemblies. Scaffolds were assembled either using crowding agents ((PEG (1-10% w/v), PVP (10% w/v), Ficoll (10% w/v), Dextran (10% w/v) or Lysozyme (10% w/v)) or monoclonal mouse antibodies (IgM) raised against human CDK5RAP2 CM2-domain (Invitrogen). The commercial antibody stock (5-10mg/mL) was diluted 1:100 in reaction buffer prior to its use. To induce CDK5RAP2 assembly using anti- CDK5RAP2 IgMs, 1μL of the diluted antibody stock was added to 9μL of the assembly reaction containing GFP::CDK5RAP2 diluted in reaction buffer. Final antibody concentration was 5-10μg/mL (5-11pM). Assembly of pre-dephosphorylated WT and S613E GFP::CDK5RAP2 scaffolds as well as reactions containing mCherry or mCherry::HSET were also induced using antibodies (120nM GFP::CDK5RAP2 final concentration). Control mCherry, full-length and ΔIDR mCherry::HSET stocks were diluted using 25mM HEPES pH7.0 prior to their addition to match salt concentrations in reaction buffer. Final concentrations of these proteins was 50nM (10μL volume).

### PLK-1 phosphorylation reactions and CDK5RAP2 phospho-mapping

PLK-1-driven expansion of GFP::CDK5RAP2 assemblies *in vitro* was achieved by incubating pre-dephosphorylated GFP::CDK5RAP2 with purified constitutively active (CA) human PLK-1 (T194D). The reaction took place at room temperature for 1 hour in reaction buffer supplemented with 20x ATP·MgCl2 (final concentration = 0.5mM). As a control we incubated CDK5RAP2 with purified kinase dead (KD) human PLK-1 (K82A) under the same conditions. Final concentrations were 120μM GFP::CDK5RAP2, 50nM PLK-1 (KD/CA), 200μM ATP, 5mM MgCl2, 50mM KCl, 25mM HEPES, pH7.0 in 10μL total volume. To identify PLK-1 phosphorylation sites in CDK5RAP2, 4.00µM of pre-dephosphorylated unlabeled CDK5RAP2 was incubated with 1µM of active human PLK- 1 (MedChem Express), 1mM ATP, 25mM MgCl2 for 1 hour at room temperature in 150mM KCl, 25mM HEPES, pH7.4 (10µL total volume). Reactions are stopped by addition of 2X SDS loading sample buffer, ran on a 4-20% protean TGX gel for 35min at 200V, and stained with a commassie-based stain InstantBlue. CDK5RAP2 bands located at the height of 215kDa were excised under sterile conditions and sent for phosphorylation PTMID. Three other CDK5RAP2 samples with no PLK-1 were submitted as non-phosphorylated controls. Samples were digested with Trypsin, Chymotrypsin and Glu-C and results were combined to maximize protein coverage (>80%). Positive phosphorylated residues were those found in the PLK-1 + ATP·MgCl2 condition and not in control reactions.

### *In vitro* aster formation

Aster assembly began by preparing a 1:5 labeled to unlabeled porcine tubulin mix on ice. To do this, we combined two tubes of 100μM commercial porcine tubulin labeled with HiLyte647 resuspended in 2.88μL of ultra-pure deionized water (5.76μL total). Labeled tubulin is then combined with 10.62μL of 137μM unlabeled porcine tubulin and 1.82μL of 100mM GTP. Final mix consists of 100μM tubulin, 10mM GTP. This mix is centrifuged (80,000rpm for 20 minutes at 4°C) using an Optima MAX-XP Ultracentrifuge in a TLA100 rotor. The top 14 microliters of the mix are transferred to a PCR tube on ice for use without disrupting precipitated tubulin. Tubulin polymerization is assessed prior to aster formation by combining 2μL of the spun tubulin mix with 1.5μL of 50% glycerol (w/v), 1μL 5XBRB80, and 0.5μL 10mM DTT. Under these conditions, numerous microtubules should spontaneously polymerize and visualized using a fluorescent microscope. In parallel, GFP::CDK5RAP2 scaffolds are prepared by combining pre-dialyzed GFP::CDK5RAP2 (WT or F75A) in reaction buffer (1μM final), 1μL of 1:100 anti-CDK5RAP2 IgMs diluted in reaction buffer (6.25–12.5 μg/mL final), 2μL of 200nM mBFP::γTuRCs in reaction buffer (50nM final), and reaction buffer up to 8µL in PCR tubes at room temperature. We can incorporate HSET into the reaction for the clustering assays by substituting 0.4μL of reaction buffer with 8μM mCherry::HSET in reaction buffer (0.4μM final). Aster reactions are also assembled in PCR tubes on ice by combining 1.5μL of 50% glycerol (w/v), 1μL 5XBRB80, 0.5μL of 10mM DTT, 1.2μL of pre-assembled CDK5RAP2 scaffolds + γTuRCs (±HSET), 0.8μL of spun tubulin mix (13µm final) and 1μL of 20X ATP·MgCl2 (666μM final) mixed with catalase (0.1mg/mL final) and glucose oxidase (1mg/mL final). All samples were then immediately transferred to a glass slide and imaged at 37°C using a TokaiHit incubation chamber. In reactions where ATP·MgCl2 was not required, equal amounts of catalase and glucose oxidase diluted in water were added instead.

### Recruitment of soluble tubulin to CDK5RAP2 scaffolds by HSET

A 100µM tubulin mixture was prepared without GTP (1:5 HiLyte647 labeled tubulin to unlabeled tubulin). To ensure removal of aggregates, the tubulin mix was spun at 80,000rpm for 20 minutes at 4°C using an Optima MAX-XP Ultracentrifuge in a TLA100 rotor. Only the top 3µL of the spun tubulin mix was used for the assays and kept on ice. 1µL of the tubulin mix was combined with 3.5µL of pre-assembled GFP- CDK5RAP2/mCherry-HSET assemblies (581nM and 593nM final concentrations respectively), 1µL of 20X ATP·MgCl2 (0.666nM ATP,16.65mM MgCl_2_ final concentration), and 0.5µL of 10mM DTT (833nM final concentration)(6µL final volume). No glycerol was added to the mix. Tubulin concentration into CDK5RAP2 assemblies was assessed by capturing far-red signal (647nm) taking 41 x 0.5µm Z-stacks using a 100X 1.35 NA silicone objective confocal microscope at 23°C. 488-nm excitation (15% laser power, 100ms exposure) followed by 561nm (10% laser power, 100ms exposure) and 647nm (10% laser power, 300ms exposure). Iris intensity was 92.3%. Images were captured using 2 × 2 binning.

### Microscopy of CDK5RAP2 scaffolds and *in vitro* asters

Upon assembly in PCR tubes scaffold reactions and aster reactions are transferred to 22×22mm microscope cover slips through gentle pipetting. Samples are then sandwiched by placing the cover slip over pre-cleaned 75×25×1mm microscope slides (VWR Vistavision) and sealed with non-toxic transparent nail polish. Then immediately transferred to a TokaiHit incubation chamber set at 37°C mounted on a Nikon Eclipse Ti2 Microscope for imaging. For clustering assays, we passivated both the cover slip and slide with mPeg-Silane the day prior imaging using a standard passivation protocol. Time-lapse images were acquired with a Yokogawa confocal scanner unit (CSU-W1), piezo Z stage, and an iXon Ultra 888 EMCCD camera (Andor), controlled by Nikon Elements software. We used a 60×1.2 NA Plan Apochromat water-immersion objective or 100X 1.35 NA silicon objective to acquire 41 × 0.5-µm Z-stacks with 100 ms exposures every two min for 30 min. 488-nm excitation (15% laser power) was recorded using 2 × 2 binning followed by 405nm excitation (10% laser power, 2×2 binning), 561nm excitation (10% laser power, 2×2 binning) and 647nm excitation (10% laser power, 1×1 binning). Iris intensity was set at 92.3%.

### Image quantification and statistical analysis

Images were analyzed using semiautomated, threshold-based particle analysis in FIJI. Data were plotted and statistical tests were performed using GraphPad prism. The sample size, measurement type, error type, and statistical test are described in the figure legends, where appropriate. Statistical significance (*) is defined as *p-value<0.05*, (****) is defined as *p-value<0.00005*, ns=not significant. Integrated fluorescence intensity is defined as area of an assembly multiplied by its mean fluorescence intensity. This value is displayed in arbitrary units (a. u.). Partition coefficient (PC) is measured by 1) identifying CDK5RA2 scaffolds using a threshold-based system (10 standard deviations above background) 2) measuring the mean fluorescence intensity of mBFP/mCherry/HiLyte-647 Alexa Fluor inside any given assembly after camera noise subtraction and 3) dividing that mean fluorescence intensity by background mean fluorescence intensity after camera noise subtraction. PC <1 indicates exclusion from GFP::CDK5RAP2 assemblies. PC = 1 indicates no selective enrichment or exclusion. PC >1 indicates selective enrichment within GFP::CDK5RAP2 assemblies. Aspect ratio is defined as the ratio between the longest and shortest diameter of a given assembly. Perfectly circular objects will have an aspect ratio of 1 while elongated objects (such as rectangles) will have aspect ratios >1.

---

**FIGURE S1.**
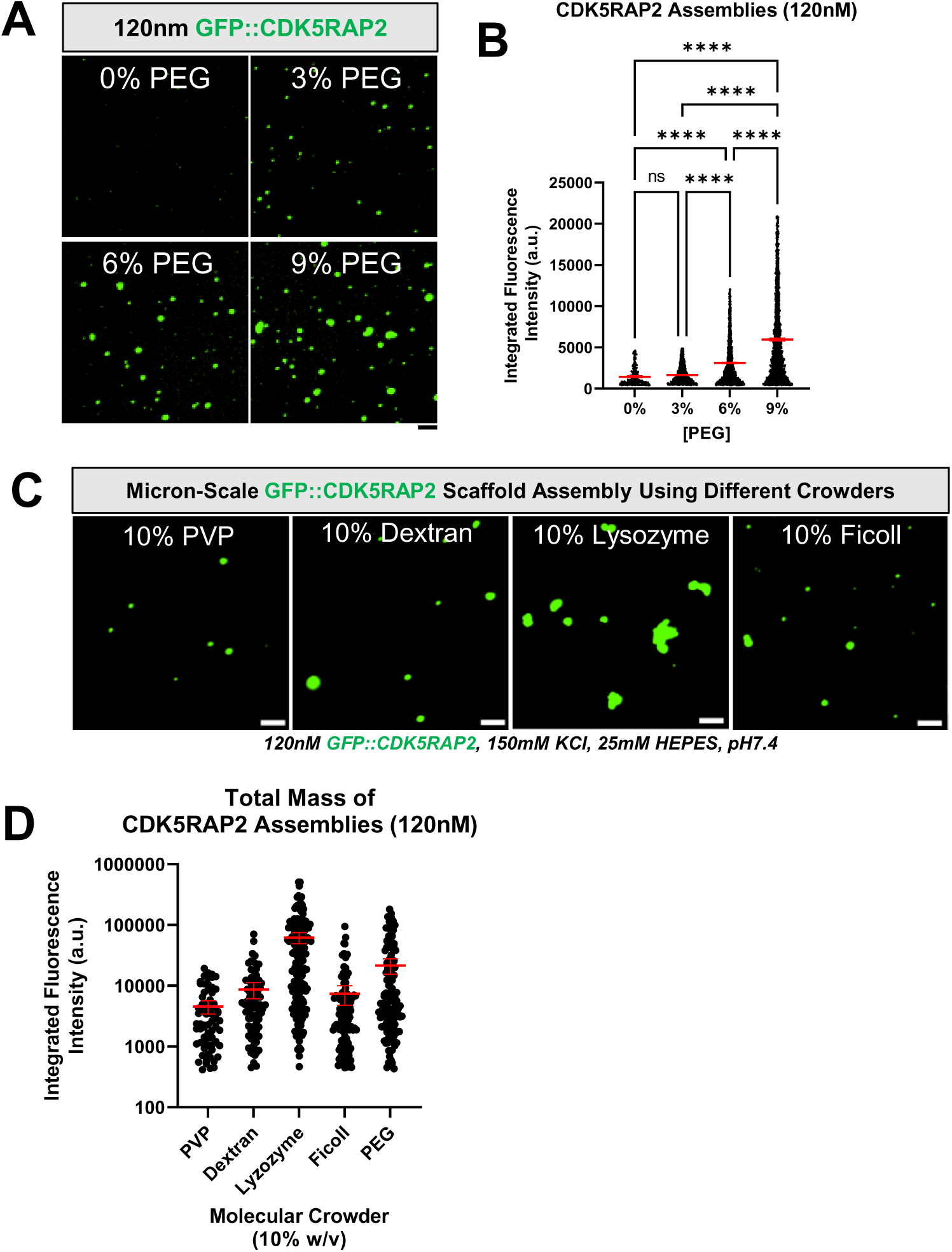
Assembly of CDK5RAP2 scaffolds using different PEG concentrations and different crowders. A) 120nM purified GFP::CDK5RAP2 combined with various PEG concentrations (0%, 3%, 6%, 9%)(w/v). Scale bar, 5μm. B) Quantification of panel 1B. Significant differences were assessed using One-way ANOVA followed by Tukey’s multiple comparisons test. Y-axis represents integrated fluorescence intensity(area x mean intensity) of CDK5RAP2 assemblies generated at various PEG concentrations. C) Micron scale assemblies of GFP::CDK5RAP2 generated using 10% (w/v) PVP, Dextran, Lyzozyme or Ficoll. Scale bar, 5μm. D) Quantification of panel 1C (mean +/- 95% C.I.; PVP (n=69 assemblies), Dextran (n=79 assemblies), Lyzozyme (n=173 assemblies), Ficoll (n=105 assemblies), PEG (n=130 assemblies)) Y-axis represents integrated fluorescence intensity (area x mean intensity) of CDK5RAP2 represented in Log10 scale.

**FIGURE S2.**
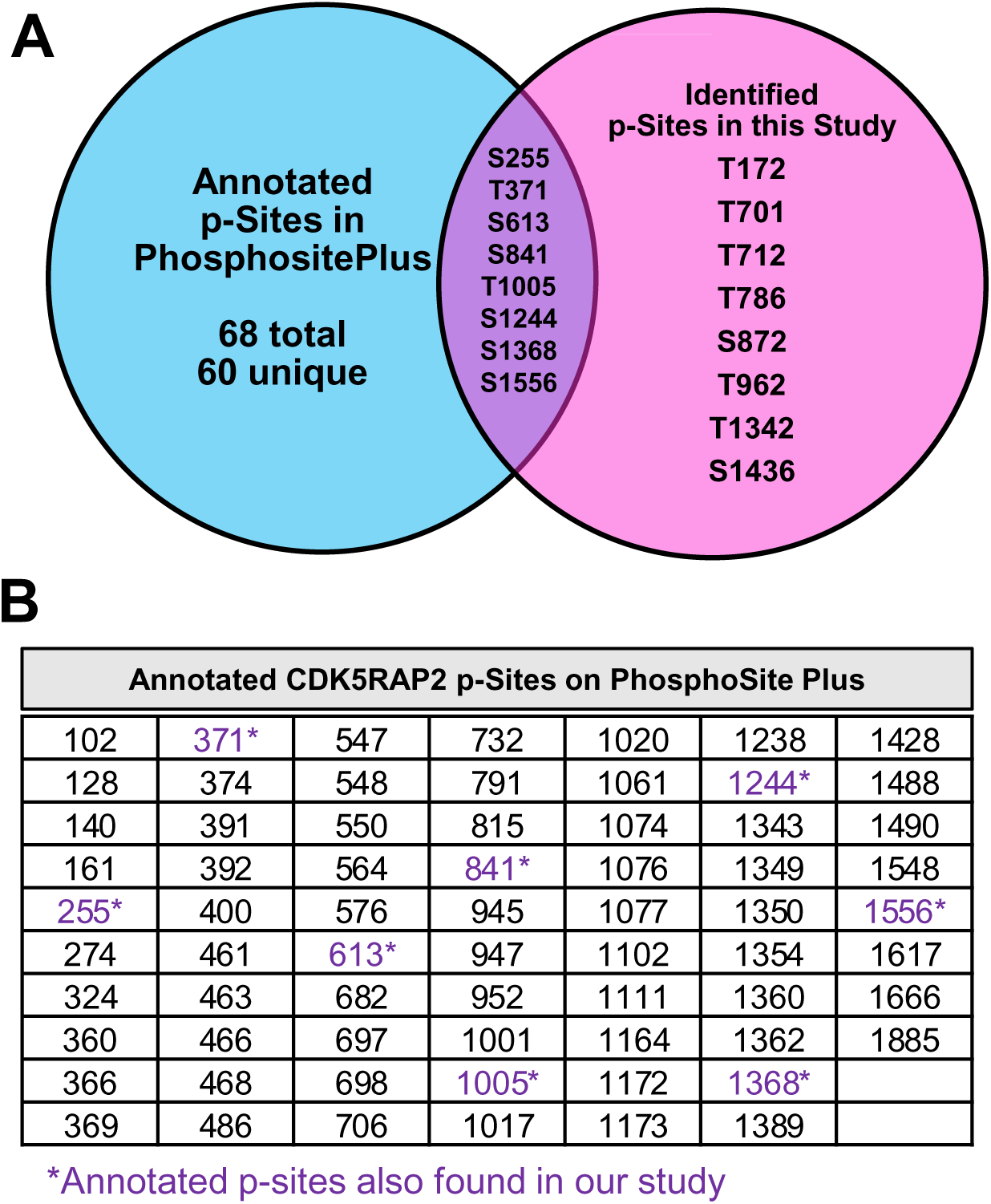
CDK5RAP2 phospho-site information A) Venn diagram indicates shared and unique p-Sites in CDK5RAP2 reported in the phospho-proteome database PhosphoSite Plus (PSP)(Blue), the phospho-sites found in our study (Pink) and the phospho-sites found in both (Purple). B) List of annotated phospho-sites in PhosphoSite Plus (Hornbeck et al., 2015). Annotated residues also found in our study are in purple.

**FIGURE S3.**
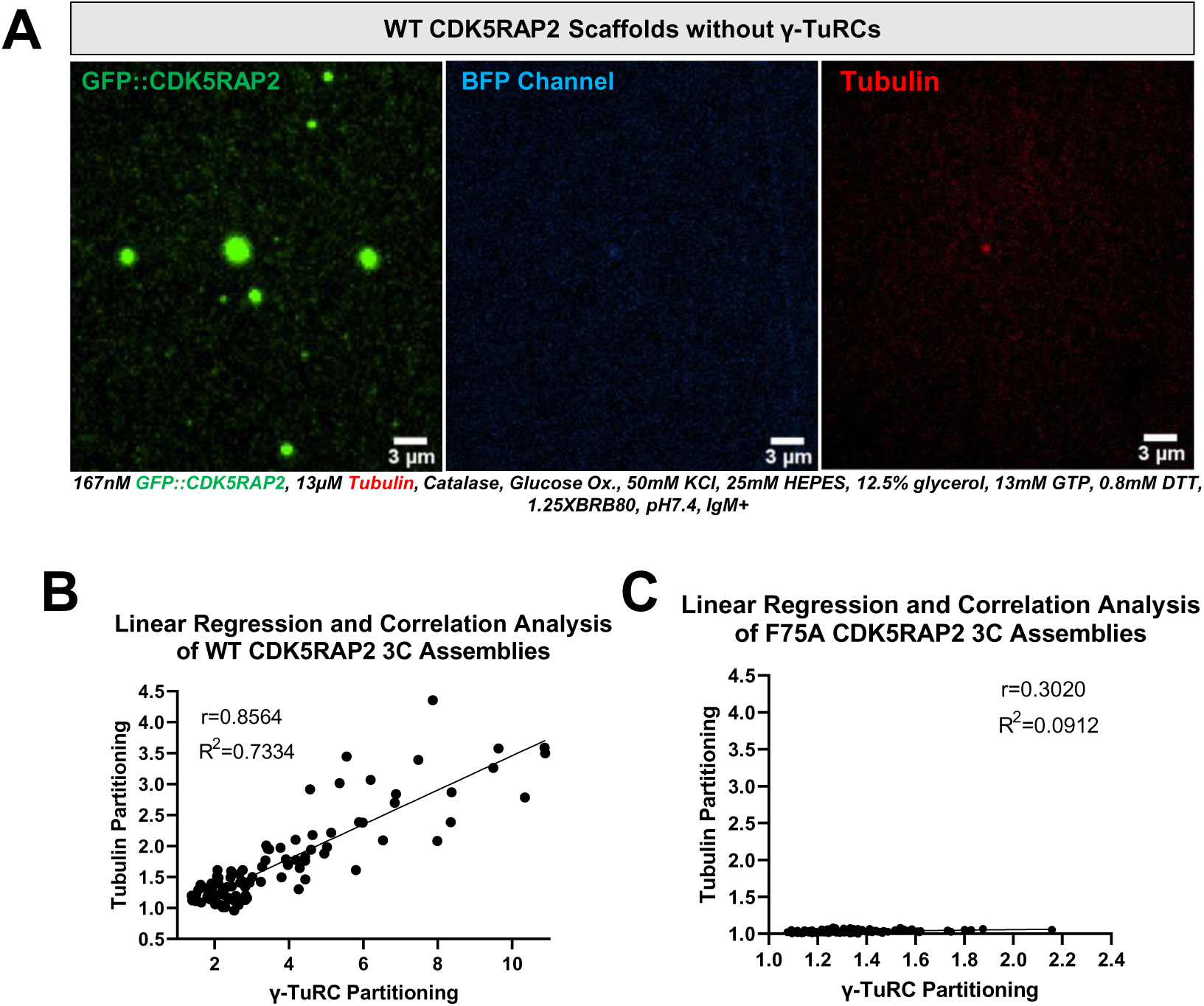
Regression analysis of 3C asters and microtubule nucleation capabilities of CDK5RAP2 alone A) 167nM GFP::CDK5RAP2(WT) scaffolds assembled using anti-CM2 IgMs in the presence of 13μM Hi-Lyte-labeled α/β tubulin mix (25mM HEPES, 50mM KCl, pH7.4). B) Linear regression and correlation analysis of CDK5RAP2(WT) assemblies. γ-TuRC partitioning (X-axis) is plotted against α/β tubulin partitioning (Y-axis) within any given CDK5RAP2 assembly. Each data point represents a CDK5RAP2 scaffold (n=95). C) Linear regression and correlation analysis of CDK5RAP2(F75A) assemblies. γ-TuRC partitioning (X-axis) is plotted against α/β tubulin partitioning (Y-axis) within any given CDK5RAP2 assembly. Each data point represents a CDK5RAP2 scaffold (n=117).

**FIGURE S4.**
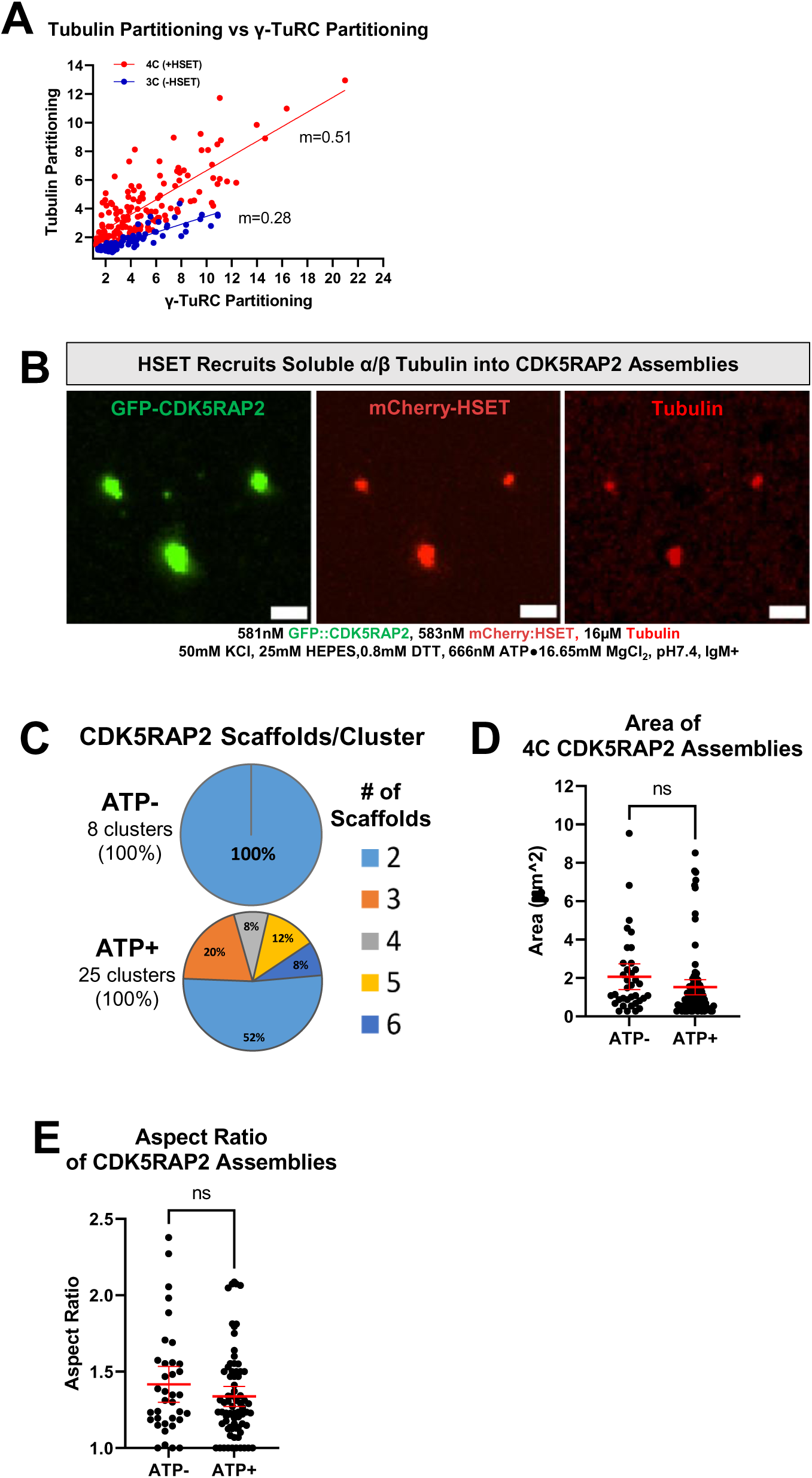
Additional analyses of CDK5RAP2 assemblies containing HSET A) Regression analysis of 4C (+HSET) vs 3C (-HSET) asters. γ-TuRC partitioning (X- axis) is plotted against α/β tubulin partitioning (Y-axis). Each data point represents a GFP-CDK5RAP2 scaffold (3C, n=95; 4C, n=133). B) mCherry-HSET can recruit α/β tubulin to GFP-CDK5RAP2 assemblies in the absence of GTP and glycerol at 23°C. Reactions contain 581nM GFP-CDK5RAP2, 583nM mcherry-HSET, 16µM Hi-Lyte-labeled α/β tubulin mix, anti-CM2 IgM. Buffer conditions are: 25mM HEPES, 50mM KCl, 666nM ATP, 16.65mM MgCl_2_ pH7.4. Scale bar, 3µm. C) Quantification of clusters and CDK5RAP2 scaffolds per cluster in ATP+/- conditions. Clusters are defined as local collections of CDK5RAP2 scaffolds connected via microtubule asters. ATP- (8 clusters from 5 images), ATP+ (25 clusters from 5 images). D) Area of GFP::CDK5RAP2 fluorescence at 4C scaffolds (mean +/- 95% C.I.; ATP, n=36 assemblies; ATP+, n=87 assemblies). Significant differences were assessed using a student t-test. E) Aspect ratio of 4C scaffolds (mean +/- 95% C.I.; ATP, n=36 assemblies; ATP+, n=87 assemblies). Significant differences were assessed using a student t-test.

**Table S1.**

| Descriptive Statistics of GFP::CDK5RAP2 Assemblies (125nm) Using Anti-CM2 IgM |  |  |  |  |  |
| --- | --- | --- | --- | --- | --- |
| N=166 | Area (µm^2) | Mean Int. | Max Int. | Circularity | Diameter (µm) |
| Minimum | 0.7510 | 3518 | 3924 | 0.7790 | 1.226 |
| Maximum | 7.323 | 31853 | 64601 | 1.000 | 3.494 |
| Range | 6.572 | 28335 | 60677 | 0.2210 | 2.268 |
| Mean | 1.875 | 8698 | 15068 | 0.9887 | 1.835 |
| Std. Deviation | 1.429 | 4397 | 10830 | 0.03598 | 0.5865 |
| Std. Error of Mean | 0.1109 | 341.2 | 840.6 | 0.002793 | 0.04552 |

**Table S2.**

| PROTEIN | JWV | pOCC Vector | Purification Tags (PTs) | Fluorescent Tags |
| --- | --- | --- | --- | --- |
| CEP215/CDK5RAP2 (FL) | 92 | pOCC29 | N-term MBP-PreScission, C-term PreScission-6xHis | N-term eGFP |
| CEP215/CDK5RAP2 (F75A) | 145 | pOCC29 | N-term MBP-PreScission, C-term PreScission-6xHis | N-term eGFP |
| CEP215/CDK5RAP2 ΔCM2 (Δ1715-1814) | 154 | pOCC29 | N-term MBP-PreScission, C-term PreScission-6xHis | N-term eGFP |
| HSET | 151 | pOCC195 | C-term PreScission-6xHis | C-term mCherry |
| HSET(ΔIDR) (Δ2-150) | 152 | pOCC195 | C-term PreScission-6xHis | C-term mCherry |
| PLK-1(T210D) | 142 | pOCC7 | C-term PreScission-6xHis | None |
| KD PLK-1 (K82A) | 144 | pOCC7 | C-term PreScission-6xHis | None |
| PROTEIN | JWB | Plasmid Name/Number | Purification Tags (PTs) | Fluorescent Tags |
| mCherry | 68 | pET AviTag His6 mCherry LIC cloning vector, Plasmid #29722 | N-term 6xHis-TEV | Itself |

## REFERENCES

1. Andersen, J.S., C.J. Wilkinson, T. Mayor, P. Mortensen, E.A. Nigg, and M. Mann. 2003. Proteomic characterization of the human centrosome by protein correlation profiling. Nature. 426:570–574.

2. Brito, C., M. Serna, P. Guerra, O. Llorca, and T. Surrey. 2024. Transition of human γ- tubulin ring complex into a closed conformation during microtubule nucleation. Science. 383:870–876.

3. Buchman, J.J., H.-C. Tseng, Y. Zhou, C.L. Frank, Z. Xie, and L.-H. Tsai. 2010. Cdk5rap2 Interacts with Pericentrin to Maintain the Neural Progenitor Pool in the Developing Neocortex. Neuron. 66:386–402.

4. Cabral, G., T. Laos, J. Dumont, and A. Dammermann. 2019. Differential Requirements for Centrioles in Mitotic Centrosome Growth and Maintenance. Developmental Cell. 50:355–366.e356.

5. Chavali, P.L., G. Chandrasekaran, A.R. Barr, P. Tátrai, C. Taylor, E.K. Papachristou, C.G. Woods, S. Chavali, and F. Gergely. 2016. A CEP215–HSET complex links centrosomes with spindle poles and drives centrosome clustering in cancer. Nature Communications. 7:11005.

6. Chen, F., J. Wu, M.K. Iwanski, D. Jurriens, A. Sandron, M. Pasolli, G. Puma, J.Z. Kromhout, C. Yang, W. Nijenhuis, L.C. Kapitein, F. Berger, and A. Akhmanova. 2022. Self-assembly of pericentriolar material in interphase cells lacking centrioles. Elife. 11.

7. Ching, Y.P., Z. Qi, and J.H. Wang. 2000. Cloning of three novel neuronal Cdk5 activator binding proteins. Gene. 242:285–294.

8. Choi, Y.-K., P. Liu, S.K. Sze, C. Dai, and R.Z. Qi. 2010. CDK5RAP2 stimulates microtubule nucleation by the γ-tubulin ring complex. Journal of Cell Biology. 191:1089–1095.

9. Conduit, Paul T., Z. Feng, Jennifer H. Richens, J. Baumbach, A. Wainman, Suruchi D. Bakshi, J. Dobbelaere, S. Johnson, Susan M. Lea, and Jordan W. Raff. 2014. The Centrosome-Specific Phosphorylation of Cnn by Polo/Plk1 Drives Cnn Scaffold Assembly and Centrosome Maturation. Developmental Cell. 28:659–669.

10. Conduit, P.T., A. Wainman, and J.W. Raff. 2015. Centrosome function and assembly in animal cells. Nat Rev Mol Cell Biol. 16:611–624.

11. Decker, M., S. Jaensch, A. Pozniakovsky, A. Zinke, K.F. O’Connell, W. Zachariae, E. Myers, and A.A. Hyman. 2011. Limiting amounts of centrosome material set centrosome size in C. elegans embryos. Curr Biol. 21:1259–1267.

12. Dobbelaere, J., F. Josué, S. Suijkerbuijk, B. Baum, N. Tapon, and J. Raff. 2008. A Genome-Wide RNAi Screen to Dissect Centriole Duplication and Centrosome Maturation in Drosophila. PLOS Biology. 6:e224.

13. Feng, Z., A. Caballe, A. Wainman, S. Johnson, A.F.M. Haensele, M.A. Cottee, P.T. Conduit, S.M. Lea, and J.W. Raff. 2017. Structural Basis for Mitotic Centrosome Assembly in Flies. Cell. 169:1078–1089 e1013.

14. Fong, K.-W., Y.-K. Choi, J.B. Rattner, and R.Z. Qi. 2008. CDK5RAP2 Is a Pericentriolar Protein That Functions in Centrosomal Attachment of the γ-Tubulin Ring Complex. Molecular Biology of the Cell. 19:115–125.

15. Fong, K.-W., S.-Y. Hau, Y.-S. Kho, Y. Jia, L. He, and R.Z. Qi. 2009. Interaction of CDK5RAP2 with EB1 to Track Growing Microtubule Tips and to Regulate Microtubule Dynamics. Molecular Biology of the Cell. 20:3660–3670.

16. Fu, J., and D.M. Glover. 2012. Structured illumination of the interface between centriole and peri-centriolar material. Open Biol. 2:120104.

17. Gell, C., C.T. Friel, B. Borgonovo, D.N. Drechsel, A.A. Hyman, and J. Howard. 2011. Purification of tubulin from porcine brain. Methods Mol Biol. 777:15–28.

18. Graser, S., Y.-D. Stierhof, and E.A. Nigg. 2007. Cep68 and Cep215 (Cdk5rap2) are required for centrosome cohesion. Journal of Cell Science. 120:4321–4331.

19. Hamill, D.R., A.F. Severson, J.C. Carter, and B. Bowerman. 2002. Centrosome maturation and mitotic spindle assembly in C. elegans require SPD-5, a protein with multiple coiled-coil domains. Dev Cell. 3:673–684.

20. Hanafusa, H., S. Kedashiro, M. Tezuka, M. Funatsu, S. Usami, F. Toyoshima, and K. Matsumoto. 2015. PLK1-dependent activation of LRRK1 regulates spindle orientation by phosphorylating CDK5RAP2. Nature Cell Biology. 17:1024–1035.

21. Haren, L., T. Stearns, and J. Lüders. 2009. Plk1-Dependent Recruitment of γ-Tubulin Complexes to Mitotic Centrosomes Involves Multiple PCM Components. PLOS ONE. 4:e5976.

22. Harrison, L.E., M. Bleiler, and C. Giardina. 2018. A look into centrosome abnormalities in colon cancer cells, how they arise and how they might be targeted therapeutically. Biochem Pharmacol. 147:1–8.

23. He, Y., Y. Shao, Z. Zhou, T. Li, Y. Gao, X. Liu, G. Yuan, G. Yang, L. Zhang, and F. Li. 2024. MORC2 regulates RBM39-mediated CDK5RAP2 alternative splicing to promote EMT and metastasis in colon cancer. Cell Death Dis. 15:530.

24. Hein, Marco Y., Nina C. Hubner, I. Poser, J. Cox, N. Nagaraj, Y. Toyoda, Igor A. Gak, I. Weisswange, J. Mansfeld, F. Buchholz, Anthony A. Hyman, and M. Mann. 2015. A Human Interactome in Three Quantitative Dimensions Organized by Stoichiometries and Abundances. Cell. 163:712–723.

25. Hentrich, C., and T. Surrey. 2010. Microtubule organization by the antagonistic mitotic motors kinesin-5 and kinesin-14. Journal of Cell Biology. 189:465–480.

26. Hornbeck, P.V., B. Zhang, B. Murray, J.M. Kornhauser, V. Latham, and E. Skrzypek. 2015. PhosphoSitePlus, 2014: mutations, PTMs and recalibrations. Nucleic Acids Res. 43:D512–520.

27. Jumper, J., R. Evans, A. Pritzel, T. Green, M. Figurnov, O. Ronneberger, K. Tunyasuvunakool, R. Bates, A. Žídek, A. Potapenko, A. Bridgland, C. Meyer, S.A.A. Kohl, A.J. Ballard, A. Cowie, B. Romera-Paredes, S. Nikolov, R. Jain, J. Adler, T. Back, S. Petersen, D. Reiman, E. Clancy, M. Zielinski, M. Steinegger, M. Pacholska, T. Berghammer, S. Bodenstein, D. Silver, O. Vinyals, A.W. Senior, K. Kavukcuoglu, P. Kohli, and D. Hassabis. 2021. Highly accurate protein structure prediction with AlphaFold. Nature. 596:583–589.

28. Kim, S., and K. Rhee. 2014. Importance of the CEP215-Pericentrin Interaction for Centrosome Maturation during Mitosis. PLOS ONE. 9:e87016.

29. Lawo, S., M. Hasegan, G.D. Gupta, and L. Pelletier. 2012. Subdiffraction imaging of centrosomes reveals higher-order organizational features of pericentriolar material. Nature Cell Biology. 14:1148–1158.

30. Lemaitre, R.P., A. Bogdanova, B. Borgonovo, J.B. Woodruff, and D.N. Drechsel. 2019. FlexiBAC: a versatile, open-source baculovirus vector system for protein expression, secretion, and proteolytic processing. BMC Biotechnol. 19:20.

31. Megraw, T.L., J.T. Sharkey, and R.S. Nowakowski. 2011. Cdk5rap2 exposes the centrosomal root of microcephaly syndromes. Trends Cell Biol. 21:470–480.

32. Mennella, V., B. Keszthelyi, K.L. McDonald, B. Chhun, F. Kan, G.C. Rogers, B. Huang, and D.A. Agard. 2012. Subdiffraction-resolution fluorescence microscopy reveals a domain of the centrosome critical for pericentriolar material organization. Nat Cell Biol. 14:1159–1168.

33. Mittasch, M., V.M. Tran, M.U. Rios, A.W. Fritsch, S.J. Enos, B. Ferreira Gomes, A. Bond, M. Kreysing, and J.B. Woodruff. 2020. Regulated changes in material properties underlie centrosome disassembly during mitotic exit. Journal of Cell Biology. 219.

34. Mountain, V., C. Simerly, L. Howard, A. Ando, G. Schatten, and D.A. Compton. 1999. The kinesin-related protein, HSET, opposes the activity of Eg5 and cross-links microtubules in the mammalian mitotic spindle. J Cell Biol. 147:351–366.

35. Nasser, H., L. Vera, M. Elmaleh-Bergès, K. Steindl, P. Letard, N. Teissier, A. Ernault, F. Guimiot, A. Afenjar, M.L. Moutard, D. Héron, Y. Alembik, M. Momtchilova, P. Milani, N. Kubis, N. Pouvreau, M. Zollino, S. Guilmin Crepon, F. Kaguelidou, P. Gressens, A. Verloes, A. Rauch, V. El Ghouzzi, S. Drunat, and S. Passemard. 2020. CDK5RAP2 primary microcephaly is associated with hypothalamic, retinal and cochlear developmental defects. J Med Genet. 57:389–399.

36. Norris, S.R., S. Jung, P. Singh, C.E. Strothman, A.L. Erwin, M.D. Ohi, M. Zanic, and R. Ohi. 2018. Microtubule minus-end aster organization is driven by processive HSET-tubulin clusters. Nature Communications. 9:2659.

37. Ohta, M., Z. Zhao, D. Wu, S. Wang, J.L. Harrison, J.S. Gómez-Cavazos, A. Desai, and K.F. Oegema. 2021. Polo-like kinase 1 independently controls microtubule-nucleating capacity and size of the centrosome. J Cell Biol. 220.

38. Pimenta-Marques, A., and M. Bettencourt-Dias. 2020. Pericentriolar material. Curr Biol. 30:R687–R689.

39. Rahimian, M., M. Askari, N. Salehi, A. Riccio, M. Jaafarinia, N. Almadani, and M. Totonchi. 2023. A novel missense variant in CDK5RAP2 associated with non-obstructive azoospermia. Taiwan J Obstet Gynecol. 62:830–837.

40. Rai, D., Y. Song, S. Hua, K. Stecker, J.L. Monster, V. Yin, R. Stucchi, Y. Xu, Y. Zhang, F. Chen, E.A. Katrukha, M. Altelaar, A.J.R. Heck, M. Wieczorek, K. Jiang, and A. Akhmanova. 2024. CAMSAPs and nucleation-promoting factors control microtubule release from γ-TuRC. Nature Cell Biology. 26:404–420.

41. Rios, M.U., M.A. Bagnucka, B.D. Ryder, B. Ferreira Gomes, N.E. Familiari, K. Yaguchi, M. Amato, W.E. Stachera, Ł.A. Joachimiak, and J.B. Woodruff. 2024. Multivalent coiled-coil interactions enable full-scale centrosome assembly and strength. Journal of Cell Biology. 223.

42. Roostalu, J., J. Rickman, C. Thomas, F. Nédélec, and T. Surrey. 2018. Determinants of Polar versus Nematic Organization in Networks of Dynamic Microtubules and Mitotic Motors. Cell. 175:796–808.e714.

43. Santamaria, A., B. Wang, S. Elowe, R. Malik, F. Zhang, M. Bauer, A. Schmidt, H.H. Silljé, R. Körner, and E.A. Nigg. 2011. The Plk1-dependent phosphoproteome of the early mitotic spindle. Mol Cell Proteomics. 10:M110.004457.

44. Schnackenberg, B.J., A. Khodjakov, C.L. Rieder, and R.E. Palazzo. 1998. The disassembly and reassembly of functional centrosomes *in vitro*. Proceedings of the National Academy of Sciences. 95:9295–9300.

45. Serna, M., F. Zimmermann, C. Vineethakumari, N. Gonzalez-Rodriguez, O. Llorca, and J. Lüders. 2024. CDK5RAP2 activates microtubule nucleator γTuRC by facilitating template formation and actin release. Developmental Cell. 59:3175–3188.e3178.

46. Sunkel, C.E., and D.M. Glover. 1988. Polo, a mitotic mutant of Drosophila displaying abnormal spindle poles. Journal of Cell Science. 89:25–38.

47. Varadi, M., S. Anyango, M. Deshpande, S. Nair, C. Natassia, G. Yordanova, D. Yuan, O. Stroe, G. Wood, A. Laydon, A. Žídek, T. Green, K. Tunyasuvunakool, S. Petersen, J. Jumper, E. Clancy, R. Green, A. Vora, M. Lutfi, M. Figurnov, A. Cowie, N. Hobbs, P. Kohli, G. Kleywegt, E. Birney, D. Hassabis, and S. Velankar. 2021. AlphaFold Protein Structure Database: massively expanding the structural coverage of protein-sequence space with high-accuracy models. Nucleic Acids Research. 50:D439–D444.

48. Varadi, M., D. Bertoni, P. Magana, U. Paramval, I. Pidruchna, M. Radhakrishnan, M. Tsenkov, S. Nair, M. Mirdita, J. Yeo, O. Kovalevskiy, K. Tunyasuvunakool, A. Laydon, A. Žídek, H. Tomlinson, D. Hariharan, J. Abrahamson, T. Green, J. Jumper, E. Birney, M. Steinegger, D. Hassabis, and S. Velankar. 2023. AlphaFold Protein Structure Database in 2024: providing structure coverage for over 214 million protein sequences. Nucleic Acids Research. 52:D368–D375.

49. Vitre, B., N. Taulet, A. Guesdon, A. Douanier, A. Dosdane, M. Cisneros, J. Maurin, S. Hettinger, C. Anguille, M. Taschner, E. Lorentzen, and B. Delaval. 2020. IFT proteins interact with HSET to promote supernumerary centrosome clustering in mitosis. EMBO Rep. 21:e49234.

50. Wang, Z., T. Wu, L. Shi, L. Zhang, W. Zheng, J.Y. Qu, R. Niu, and R.Z. Qi. 2010. Conserved motif of CDK5RAP2 mediates its localization to centrosomes and the Golgi complex. J Biol Chem. 285:22658–22665.

51. Watanabe, S., F. Meitinger, A.K. Shiau, K. Oegema, and A. Desai. 2020. Centriole-independent mitotic spindle assembly relies on the PCNT–CDK5RAP2 pericentriolar matrix. Journal of Cell Biology. 219.

52. Wong, S.S., Z.M. Wilmott, S. Saurya, I. Alvarez-Rodrigo, F.Y. Zhou, K.Y. Chau, A. Goriely, and J.W. Raff. 2022. Centrioles generate a local pulse of Polo/PLK1 activity to initiate mitotic centrosome assembly. Embo j. 41:e110891.

53. Woodruff, J.B., B. Ferreira Gomes, P.O. Widlund, J. Mahamid, A. Honigmann, and A.A. Hyman. 2017. The Centrosome Is a Selective Condensate that Nucleates Microtubules by Concentrating Tubulin. Cell. 169:1066–1077.e1010.

54. Woodruff, J.B., O. Wueseke, V. Viscardi, J. Mahamid, S.D. Ochoa, J. Bunkenborg, P.O. Widlund, A. Pozniakovsky, E. Zanin, S. Bahmanyar, A. Zinke, S.H. Hong, M. Decker, W. Baumeister, J.S. Andersen, K. Oegema, and A.A. Hyman. 2015. Centrosomes. Regulated assembly of a supramolecular centrosome scaffold in vitro. Science. 348:808–812.

55. Wueseke, O., D. Zwicker, A. Schwager, Y.L. Wong, K. Oegema, F. Jülicher, A.A. Hyman, and J.B. Woodruff. 2016. Polo-like kinase phosphorylation determines Caenorhabditis elegans centrosome size and density by biasing SPD-5 toward an assembly-competent conformation. Biol Open. 5:1431–1440.

56. Xu, Y., H. Muñoz-Hernández, R. Krutyhołowa, F. Marxer, F. Cetin, and M. Wieczorek. 2024. Partial closure of the γ-tubulin ring complex by CDK5RAP2 activates microtubule nucleation. Developmental Cell. 59:3161–3174.e3115.

57. Yigit, G., K.E. Brown, H. Kayserili, E. Pohl, A. Caliebe, D. Zahnleiter, E. Rosser, N. Bögershausen, Z.O. Uyguner, U. Altunoglu, G. Nürnberg, P. Nürnberg, A. Rauch, Y. Li, C.T. Thiel, and B. Wollnik. 2015. Mutations in CDK5RAP2 cause Seckel syndrome. Mol Genet Genomic Med. 3:467–480.

